# Fungal community assembly in soils and roots under plant invasion and nitrogen deposition

**DOI:** 10.1101/416818

**Authors:** Michala L. Phillips, S□ren E. Weber, Lela V. Andrews, Emma L. Aronson, Michael F. Allen, Edith B. Allen

## Abstract

Abstract Fungal community composition in the Anthropocene is driven by rapid changes in environmental conditions caused by human activities. This study examines the relative importance of two global change drivers – atmospheric nitrogen (N) deposition and annual grass invasion – on structuring fungal communities in a California chaparral ecosystem, with emphasis on arbuscular mycorrhizal fungi. We used molecular markers, functional groupings, generalized linear statistics and joint distribution modeling, to examine how environmental variables structure taxonomic and functional composition of fungal communities. Invasion of a chaparral ecosystem decreased richness and relative abundance of non-AMF symbionts and rhizophilic AMF (e.g. Glomeraceae) as well as the proportion of edaphophilic AMF (e.g. Gigasporaceae). We found increased richness and the proportion of rhizophilic and edaphophilic AMF with increasing soil NO_3_. Our findings suggest that invasive persistence may decrease the presence of multiple soil symbionts that native species depend on for pathogen protection and increased access to soil resources.

## Introduction

Soil fungal community composition responds strongly to drivers of global change such as non-native plant invasions and atmospheric nitrogen (N) deposition (Egerton-Warburton & Allen 2000; Amend *et al.* 2015). The U.S. southwest is experiencing high rates of invasion from Mediterranean annual grasses facilitated by increased N deposition (Ashbacher & Cleland 2015; Fenn *et al.* 2010). Decreases in plant diversity following invasion alter the composition and function of soil fungi via changes in litter inputs and symbiotic relationships (Wolfe & Klironomos 2005; Reinhart & Callaway 2006; Inderjit & van der Putten 2010). N deposition is also altering fungal composition both directly through shifts in nutrient availability and indirectly via shifts in plant community composition. While vegetation responses to invasion and N deposition have been examined (Rao & Allen 2010; Valliere *et al.* 2017), relatively little is known about soil fungal responses, despite recent efforts (Egerton-Warburton & Allen 2000; Egerton-Warburton *et al.* 2001; Egerton-Warburton, Johnson & Allen 2007; Amend *et al.* 2015).

Many fungal functional groups may respond to drivers of global change, including arbuscular mycorrhizal fungi (AMF), ectomycorrhizal fungi (EMF), saprotrophs and pathogens. AMF are plant mutualists, providing host plants with resources (nutrients and water) in exchange for photosynthetically derived carbon. N deposition and invasion of non-native plant species have the potential to shift the structure and function of both AMF and broader fungal communities. N deposition can lead to soil eutrophication, which has the potential to reduce the dependence of host-plants on AMF for nutrient uptake (Treseder & Allen 2002; Egerton-Warburton, Johnson & Allen 2007). Additionally, some invasive plants exhibit relatively low AMF dependence which could decrease the presence of AMF (Busby *et al.* 2013, 2011; Hawkes *et al.* 2006). Molecular advances have facilitated the discovery of substantial diversity within AMF. Yet, without determining the functional significance of specific AMF taxa, it is challenging to infer the ecological importance of shifts in taxa abundance (Peay 2014).

The composition of AMF may be altered by invasive annual grasses from the Mediterranean that replace shrub communities (e.g. chaparral) in southern California (Egerton-Warburton & Allen 2000). The mechanism for this shift in species composition may be related to host-specificity of AMF (Hausmann & Hawkes 2009; Sikes *et al.* 2009), which could result in differences in community composition and function between invasive and native host plants. Fast-growing AMF taxa may preferentially colonize species with earlier root activity and more fibrous root structures that are well suited for rapid nutrient uptake, such as invasive grasses (Hooper & Vitousek 1998). Increased presence of intra-radical hyphae produced by these AMF taxa confer pathogen protection to vulnerable fibrous roots (Maherali & Klironomos 2007; Sikes *et al.* 2010). Abundant fast-growing AMF taxa in the roots of invasive grasses may create a positive feedback loop and promote grass invasion. On the other hand, woody plant species such as native shrubs with slower growth rates and coarser root morphologies may be more dependent upon slower growing AMF taxa with their capacity for nutrient uptake via long extraradical hyphae (Allen *et al.* 2003; Maherali & Klironomos 2007; Hart & Reader 2002). Release from fungal pathogens could also promote the establishment of invasive plants (Mitchell & Power 2003; Kardol *et al.* 2007; Van Grunsven *et al.* 2007; Reinhart *et al.* 2010), though pathogen release is less important in disturbed systems (Müller *et al.* 2016). In resource-poor environments where plants are heavily dependent on mycorrhizal relationships, disruptions of these mutualistic networks through invasion can promote the establishment and persistence of invasive plants (Richardson *et al.* 2000; Callaway *et al.* 2008; Busby *et al.* 2013).

AMF associations are not affected by their host plants alone, but also directly and indirectly by soil properties. Previous work has shown interactive effects of nitrogen (N) and phosphorus (P) on AMF taxa, such that in P-rich soil (lower N:P ratio) nitrogen fertilization decreases AMF productivity and diversity (Treseder & Allen 2002; Egerton-Warburton, Johnson & Allen 2007). At P-limited sites, fertilization often increases AMF productivity and diversity (Treseder & Allen 2002; Egerton-Warburton, Johnson & Allen 2007). However, as nutrient availability increases, it is likely that host plants will depend less on AMF taxa that produce extraradical hyphae for nutrient uptake (Sikes *et al.* 2010). Invasion by exotic annual plants has been linked to the rise in N deposition in southern California (Rao & Allen 2010; Valliere *et al.* 2017). Therefore, invasion and N deposition may synergistically decrease the diversity and abundance of slower growing AMF families.

AMF have been previously described by AMF functional groups as early and late successional delineated by spore size (e.g. Allen *et al.* 2003). Alternatively, the guild approach outlined in Weber *et al.* (2018, this issue), organizes AMF families by patterns of biomass allocation (Table 1), synthesized from previous studies (Powell *et al.* 2009; Hart & Reader 2002; Varela-Cervero *et al.* 2015; Varela-Cervero *et al.* 2016a; Varela-Cervero *et al.* 2016b). Briefly, this approach classifies AMF families with high allocation to extradical hyphae as ‘edaphophilic,’ those with high allocation to root colonization as ‘rhizophilic,’ and those with lower allocation to either root colonization or soil hyphae than the edaphophilic or rhizophilic guilds as ‘ancestral.’ Families in the edaphophilic guild are known to improve plant nutrient uptake, whereas families in the rhizophilic guild may protect host plant roots from pathogen colonization (Sikes *et al.* 2010, Treseder *et al.* 2018).

**Table 1:**
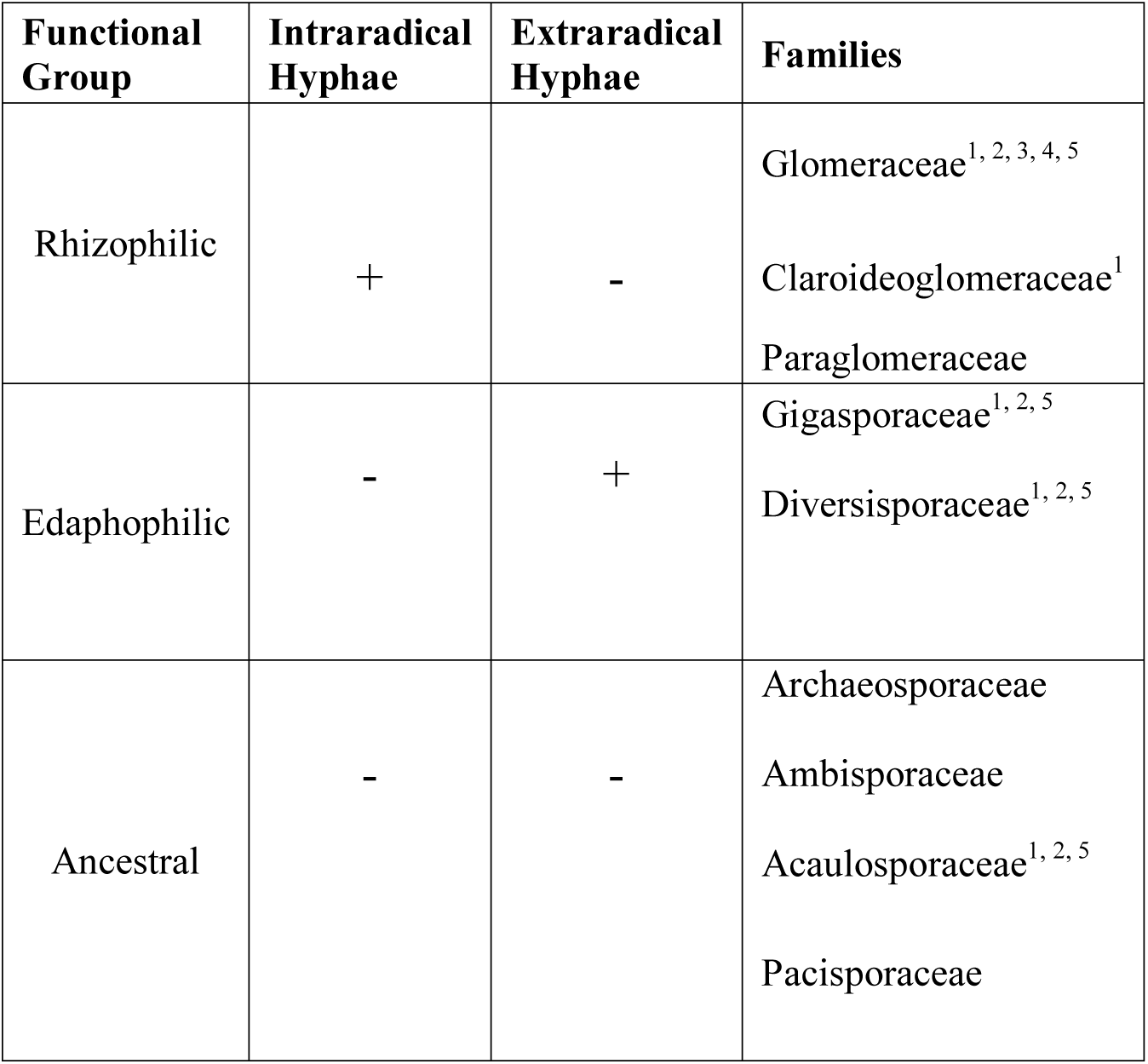
Description of AMF Functional Groups adapted from (Weber *et al*., 2017).

In this study, we focus on AMF, but also assess changes in other fungal functional groups including saprotrophs, pathogens and non-AMF symbionts, as these functional groups interact with AMF and are also affected by the same global changes drivers (Amend *et al*. 2015). We hypothesize that (1) native shrub roots will host relatively more edaphophilic AMF, whereas invasive grass roots will host relatively more rhizophilic AMF; (2) invasive grass roots will harbor fewer pathogens than native shrubs; and, (3) elevated soil N concentrations will reduce the richness and relative abundance of edaphophilic AMF taxa. We test these hypotheses within both guild and broader taxonomic frameworks, using high-throughput sequencing coupled with generalized linear models and joint taxa distribution models to understand the importance of multiple environmental variables in structuring fungal communities.

## Methods

### Site Description

We sampled from two chaparral communities in southern California, the San Dimas Experimental Forest (SDEF) and Emerson Oaks Reserve (EOR), both with granitic parent material and coarse sandy loam soils. San Dimas Experimental Forest is in the San Gabriel Mountains (34 12’ N, 117 46’ W, 50 km east of Los Angeles), at 830 m above sea level. A small portion of SDEF (∼100 ha) was purposely converted from native chaparral to grassland in the 1960s to study the relationship between ecohydrology and community type (Dunn *et al*. 1988). EOR is in Temecula Valley (33 28’ N, 117 2’ W,) 500 m in elevation. We sampled in both a grassy patch ∼1 ha where shrubs had been cleared before the 1980s and in surrounding mature chaparral. Both sites burned in wildfires within the past 20 years (SDEF – 2003, EOR – 2004), but we sampled in both areas where chaparral had recovered, and areas where exotic grasslands persisted. Because of SDEF’s proximity to Los Angeles, it receives a large amount of atmospheric N deposition (> 19 kg N ha ^-1^ yr^-1^, Fenn *et al.* 2010). EOR receives much less atmospheric N deposition (∼6 kg N ha ^-1^ yr^-1^, Fenn *et al.* 2010).

### Host plants

In March 2016, we sampled roots and bulk soils at both sites underneath individuals (n=6) of the dominant native chaparral shrub, *Adenostoma fasciculatum*. *A. fasciculatum* is a dominant shrub species in chaparral which forms several types of root-fungal associations, primarily with AMF, but also with ectomycorrhizal fungi and dark-septate fungi (Allen et al., 1999). We sampled the dominant invasive grass species (n = 6) at each site (*Bromus diandrus* at EOR and *Avena fatua* at SDEF). At each site we sampled from adjacent stands (>5 meters but <10 meters apart) of invasive and native vegetation. Sample size analysis indicated that >95% of fungal richness was likely captured with six samples (‘vegan’ package, Oksanen *et. al*, 2017).

### Soil Sampling

Soils cores were collected at ∼10 cm depth from the base of each individual plant. Roots were washed thoroughly with DI water and soils were sieved using a 2 mm mesh that was sterilized with 70% ethanol between samples. Samples were frozen at -20 °C until analyzed. Each soil sample was analyzed for pH in a DI water slurry, for KCl-extractable NH_4_ and NO_3_ (University of California Davis Analytical Laboratory), and for bicarbonate-extractable P (USDA-ARS Soils Laboratory, Reno, NV). Soil characteristics by site and host plant type are summarized in Tables 2 and 3.

**Table 2:** Soil characteristics for each site. Values shown are mean of all samples with standard error in parentheses.

| Site | pH | NH4<br>(ppm) | NO3<br>(ppm) | P (ppm) |
| --- | --- | --- | --- | --- |
| EOR | 6.69<br>(0.05) | 1.51<br>(0.07) | 2.94<br>(0.60) | 11.85<br>(0.64) |
| SDEF | 6.09<br>(0.08) | 1.76<br>(0.27) | 12.05<br>(1.89) | 7.21<br>(0.57) |

**Table 3:**
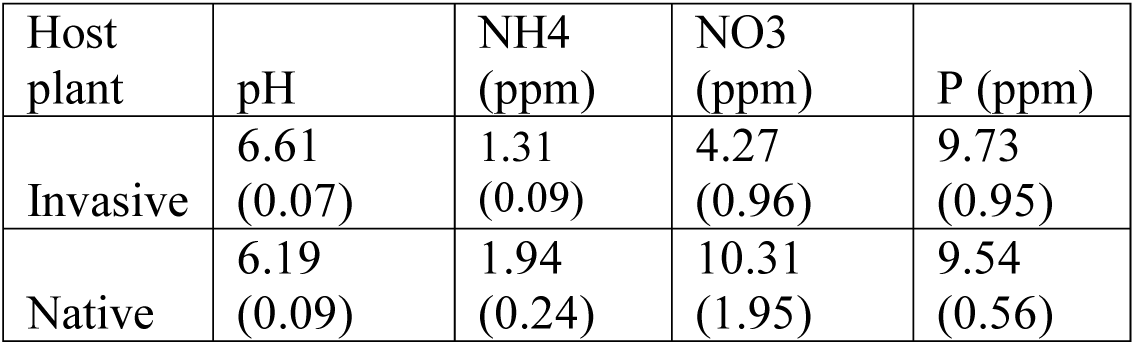
Soil characteristics for each host plant. Values shown are mean of all samples with standard error in parentheses.

We extracted DNA from soils (∼0.25g/sample) and roots (∼0.15g/sample) using the Powerlyzer PowerSoil DNA Isolation Kit per manufacturer’s protocol (Mo Bio Laboratories, Carlsbad California), with a modified heated lysis step at 65°C for twenty minutes, before homogenization (Rubin *et al*. 2014). Samples were kept frozen in a -20 °C freezer and transported on dry ice to the NAU Environmental Genetics and Genomics Laboratory (EnGGen) at Northern Arizona University. Samples were further purified from residual contaminants by the PEG-bead protocol described by Rohland & Reich 2012. DNA concentrations were determined by PicoGreen (Molecular Probes Inc., Eugene OR, USA) fluorescence and standardized to ∼10 ng/µL.

### Percent colonization

To assess fungal colonization, roots remaining after DNA extraction were washed from soil, cleared overnight in 2.5 % KOH, acidified in 1% HCl, and stained in 0.05% trypan blue (Kormanik & McGraw 1982; Koske & Gemma 1989). We estimated percent colonization using a modified magnified intersection method (McGonigle *et al.* 1990). Roots were mounted in PVLG on microscope slides and 60 intercepts per replicate were observed at 200× magnification. We examined root fragments for AMF hyphae, arbuscules, vesicles, as well as hyphae, reproductive structures of non-AM fungi, and EMF mantles and Hartig nets.

To test for differences in colonization between invasive and native hosts, five linear models were fit to percent colonization data using structures listed above as response variables and host plant, site, and host plant by site as the predictor variables. ANOVA was used to assess variable significance. All statistical analyses were performed in R version 3.2.1 (R version 3.2.1; R Core Team 2017).

### Library construction and sequencing

Samples were amplified by polymerase chain reactions (PCR) for the ribosomal small subunit (SSU) region using the Glomeromycotina-specific AML2 and the universal eukaryote WANDA primer set (Lee *et al.* 2008; Dumbrell *et al*. 2011) and for the internal transcribed spacer 2 (ITS2) region using the universal fungal primers 5.8SFun and ITS4Fun (Taylor *et al*. 2016) in preparation for high-throughput sequencing of the resulting amplicon pools. Library construction was conducted in a two-step procedure as in Berry *et al.* (2011). First-round amplifications were carried out in triplicate with three separate template dilutions (10 ng, 1 ng, or 0.1 ng template DNA), and with primers possessing universal tails synthesized 5’ to the locus specific sequences (Alvarado *et al.* 2017, in press). Besides template DNA, reactions contained 0.1 U/µL Phusion HotStart II DNA polymerase (Thermo Fisher Scientific, Waltham, MA), 1X Phusion HF Buffer (Thermo Fisher Scientific), 200 µM dNTPs (Phenix Research, Candler, NC), and 3.0 mM MgCl_2_. Thermal cycler conditions were as follows: 2 min at 95 °C; 35 cycles of 30 seconds at 95 °C, 30 seconds at 55 °C, 4 minutes at 60 °C; then refrigerate at 10 °C. Triplicate reaction products for each sample were pooled by combining 4 µL from each, and 2 µL was used to check results on a 1% agarose gel. Products were purified by the PEG-bead cleanup and eluted in 100 µL Tris-Cl pH 8.0. 1 µL of purified, diluted product was used as template in a second, indexing PCR reaction, using primers with sequences matching the universal tails at the 3’ end, and matching Illumina MiSeq flowcell sequences at the 5’ end. Conditions for tailing reactions were identical to the first round except that we used 100 nM of each indexing primer, only one reaction was conducted per sample and only 15 total cycles were performed. We used 2 µL to check results on an agarose gel, purified by the PEG-bead cleanup, quantified by PicoGreen fluorescence, and equal masses for every sample were combined into a final sample pool using an automated liquid handling system (PerkinElmer, Waltham, MA). We further concentrated the resulting pool with the PEG-bead protocol, quantified it by qPCR and average fragment sizes were estimated using a Bioanalyzer 2100 (Agilent Technologies, Santa Clara, CA) prior to sequencing. Sequencing was carried out on a MiSeq Desktop Sequencer (Illumina Inc, San Diego, CA) running in paired end 2x300 mode.

### Bioinformatics

We used cutadapt (Martin 2011) to filter sequences for locus-specific primer sequences and smalt (http://www.sanger.ac.uk/science/tools/smalt-0) to remove residual PhiX contamination, the viral genome used as a control sequence on Illumina Platforms. For the ITS loci, we joined paired-end reads with ea-utils (Aronesty 2011) and checked joined-sequence quality with FastQC (Andrews S. 2010). FastQC: a quality control tool for high throughput sequence data; available online at: http://www.bioinformatics.babraham.ac.uk/projects/fastqc). For the SSU loci, we used the forward read and checked quality with FastQC (Andrews S. 2010). Demultiplexing was performed in QIIME 1.9.1 (Caporaso *et al.* 2010) with the `split_libraries_fastq.py` command using a phred score of 20 (q = 19), allowing zero low-quality base calls (r = 0), and retaining reads only if they possess 95% of initial sequence length following quality truncation (p = 0.95). We screened for chimeras using VSEARCH (Rognes *et al.* 2016) in uchime_denovo mode for SSU and uchime_ref mode against the UNITE-based fungal chimera dataset for ITS (Nilsson *et al.* 2015). For ITS2, fungal sequences were extracted using ITSx (Bengtsson-Palme *et al.* 2013). We picked OTUs using Swarm (Mahe *et al.* 2014) with a resolution of *d4*. A resolution of *d4*, a local clustering threshold, collapses sequences with no more than 4 differences into a single representative OTU, given our quality filtering threshold of q20 and the trimmed length of amplicon sequences. Taxonomy was assigned using BLAST, with an e-value of 0.001 (Altschul *et al.* 1990) against the UNITE ITS reference database (Kõljalg *et al.* 2013) and MaarjAM database for SSU (Öpik *et al.* 2010). Reference databases were truncated prior to analysis to include only the region of interest to avoid any spurious results. For both loci, we normalized OTUs using cumulative sum scaling (CSS-normalization) in the metagenomeSeq package of Bioconductor (Paulson *et al.* 2013) in R prior to further analyses (R Core Team 2017). CSS normalization attempts to avoid biases in marker gene surveys due to uneven sequencing depth. Read counts are rescaled against a quantile determined by assessing count deviations of each sample as compared to the distribution of counts across all other samples (Paulson *et al.* 2013). Raw and CSS-normalized OTU tables are available through Mendeley Data at http://dx.doi.org/10.17632/ppmfn3rh7r.1 (Phillips, 2018).

### Functional group assignment

To examine responses of the general fungal community (ITS2), we assigned OTUs to functional groups using the online application FUNguild (“http://www.stbates.org/guilds/app.php”, Nguyen *et al.* 2016). After processing OTUs through FUNguild, we removed Glomeromycotina from the symbiont group to remove redundancy of ITS2 and SSU sequences. The remaining non-AMF symbionts include EMF. EMF occurrence was low in both native and invasive samples, so we did not analyze them separately. To simplify, FUNguild functional groups ‘pathotrophs’, ‘pathotroph-saprotrophs’ and ‘pathotroph-symbiotrophs’ were assigned to the pathogen group; and ‘saprotrophs’ and ‘saprotroph-pathotroph’ to the saprotroph group. We kept only FUNguild assignments that were at the confidence level of ‘highly probable’ and ‘probable, removing all taxa that were at the confidence level of ‘possible’ for these analyses. We retain saprotrophic FUNguild assignments in roots under the assumption that these saprotrophs may be opportunistically parasitizing plant roots, as recent research uncovers the potential for fungi to occupy multiple niches (Glynou *et al.* 2017; Selosse *et al.* 2018). With these constraints, FUNGuild was able to assign function to 585 OTUs (62%) of 940 ITS2 OTUs.

For the SSU locus, 181 OTUs (65%) out of 277 were assigned taxonomy by using BLAST against the MaarjAM database. To interpret responses of the AMF community (SSU) we assigned families of Glomeromycotina to AMF functional groups: rhizophilic, edaphophilic and ancestral using AMF resource allocation patterns defined in previous studies (Table 1). Families that did not fall into rhizophilic or edaphophilic groups were placed in the ancestral group (Table 1). We did not include sequences reportedly identified as *Geosiphon pyriformis*, of which there were only two observations, in any of the functional groups.

### Beta Diversity

For each locus, we visualized beta-diversity using non-metric multidimensional scaling (NMDS) of the Bray-Curtis distances. The NMDS was visualized in R (R version 3.2.1; R Core Team 2017) using the ggplot2 package (Wickham 2009) and the ‘stat_ellipse’ function with 95% confidence intervals. We tested for differences in overall general fungal (ITS2) and AMF (SSU) community composition across treatments by performing permutational multivariate ANOVA (PERMANOVA) for each locus using the ‘adonis’ function in the ‘vegan’ R package (999 permutations; Oksanen et al. 2017). Host plant, site, type (root or soil), pH, NO_3_, NH_4,_ and P were used as the predictor variables. For the SSU locus, we could not include pH, NO_3_, NH_4_ and P in the PERMANOVA because the multivariate homogeneity of groups dispersion was not met. For the ITS2 locus, we could include all variables as the homogeneity of groups dispersion was met for every predictor variable. For both loci, these analyses were completed on the full datasets before filtering for functional group assignment.

### Generalized linear models

We used generalized linear models (GLMs) to test our hypotheses about fungal functional group responses to invasion and elevated soil N concentrations. We built GLMs using the ‘glm’ function in the MASS package in R (Venables and Ripley et. al., 2002). We fit models using gaussian, negative binomial, poisson and log normal distributions where appropriate, determined with the ‘qqp’ function in the MASS package to visually assess probability distribution fit. We used the ‘stepAIC’ function from the MASS package to further select these models for parsimony (Venables & Ripley *et. al.* 2002). We used separate models for roots and soils by functional group richness and abundance of each locus, resulting in twenty-four models.

### Joint taxa distribution modeling

To understand how environmental variables structure AMF relative taxonomic abundance, we analyzed read abundance data (Paulson *et al.* 2013) using joint distribution models following the Hierarchical Modeling of Species Communities approach (‘HMSC’ R package) as outlined in Ovaskainen *et al.* (2017). The HMSC approach uses a hierarchical Bayesian structure to fit a joint distribution model to presence/absence or abundance data of taxa from diverse communities.

We built and evaluated models examining responses of AMF read abundance for roots and soils of the SSU locus at the family level, resulting in two models. We performed 200,000 Marcov chain Monte Carlo (MCMC) iterations of each model, of which the first half was discarded, and the remaining 100,000 were further thinned, resulting in 1,000 posterior samples. We used flat priors and sampled the posterior distribution using the Gibbs sampler with a Gaussian distribution. Both models included the same environmental predictors: host plant, site, pH, NH_4_, NO_3_, and P. We considered environmental predictors as fixed effects and individual sample as a random effect. We checked for model convergence by visually assessing the MCMC trace plots. We used the posterior distributions of each predictor and calculated the probability that it was different from zero. We considered parameters “significant” when their posterior probabilities had at least a 90% probability of being different from zero (p = 0.1). We used the ‘variPart’ function in the HMSC package to calculate the relative proportion of the total model variance that is attributable to each of the fixed and random effects (Blanchet & Tikhonov 2016). This allows us to assess the explanatory power of our models, while also understanding how much variation in family abundance can be explained by each of our environmental variables as well as random processes.

## Results

### Percent colonization

Roots of invasive annual grasses had higher colonization by AM and non-AM hyphae than native shrub roots (72% ± 4 (SD) invasive and 5% ± 33 native, P = 0.003, and 56% ± 38 and 8% ±7, P = 0.023, respectively). Rates of AMF hyphal colonization in roots were higher in both native and invasive host plants at SDEF than at EOR (55% ± 35 vs. 13% ± 11). The colonization of arbuscules (0% in native and 1% in invasive roots) was too low to analyze statistically, though we did observe more vesicles in invasive roots than in native roots (11% and 2%, respectively; P = 0.002). We did not observe ectomycorrhizal fungal (EMF) colonization in *A. fasciculatum* roots.

### SSU sequences (AMF)

We observed a total of 277 OTUs, 181 of which were assigned taxonomy after performing BLAST against the MaarjAM database. For sequences with assigned taxonomies, we observed a mean of 335±121 (SD) reads, and 52 ± 16 OTUs, per sample. These OTUs belonged to 3 orders, 10 families and 9 genera within Glomeromycotina. We observed the following 9 genera: *Glomus, Acaulospora, Archaeospora, Paraglomus, Scutellospora, Claroideoglomus, Geosiphon, Ambispora,* and *Redeckera*. Of those genera, only 2 OTU’s were identified as *Geosiphon pyriformis* which we removed from subsequent analyses, because it did not fall into any AMF functional grouping. Family abundances can be found in table S2. We placed these OTUs into three functional guilds described earlier (Table 1). Of these guilds, the most common were rhizophilic AMF (264 ± 105 reads and 39 ± 12 OTUs per sample), followed by edaphophilic families (50 ± 29 reads and 8 ± 3 OTUs per sample) with ancestral AMF being the least common (39 ±20 reads and 16 ± 6 OTUs per sample).

### ITS2 sequences (general fungal community)

We observed a mean ± SD of 661 ± 277 reads and 125 ± 50 OTUs per sample. These OTUs belonged to 7 phyla, 21 classes, 40 orders, 79 families and 149 genera. The most abundant phylum in the roots was Ascomycota with 442 ± 203 reads and 84 ± 32 OTUs per sample, followed by Basidiomycota with 182 ± 104 reads and 33 ± 18 OTUs. Saprotrophs were the most common (189 ± 219 reads and 36 ± 42 OTUs per sample), followed by pathogens (65 ± 64 reads and 13 ± 11 OTUs per sample) and non-AMF symbionts (62 ± 65 reads and 11 ± 8 OTUs per sample). Once we removed AMF to avoid overlap between our datasets, the remaining fungal symbionts consisted of 11 families, 11 genera, and 20 species. Of the 11 families, seven families – Inocybaceae, Tricholomataceae, Pyronemataceae, Sclerodermataceae, Helvellaceae, Rhizopogonaceae and Paxillaceae – contain EMF species. Four families – Collemataceae, Teloschitaceae, Lobariaceae, Lecideaceae – contain lichenized fungal species.

### Beta Diversity

AMF beta diversity differed by site (R^2^ = 0.04, P = 0.02, Figure 1a). Host plant, sample type (root or soil) and their interaction did not significantly structure AMF beta diversity (R^2^ = 0.01 and 0.02; P = 0.9 and 0.6, respectively). Beta diversity of the general fungal community was significantly structured by host plant (R^2^ = 0.04, P = 0.01, Figure 2a) and the interaction between host plant and sample type (R^2^ = 0.03, P = 0.04, Figure 2b).

**Figure 1:**
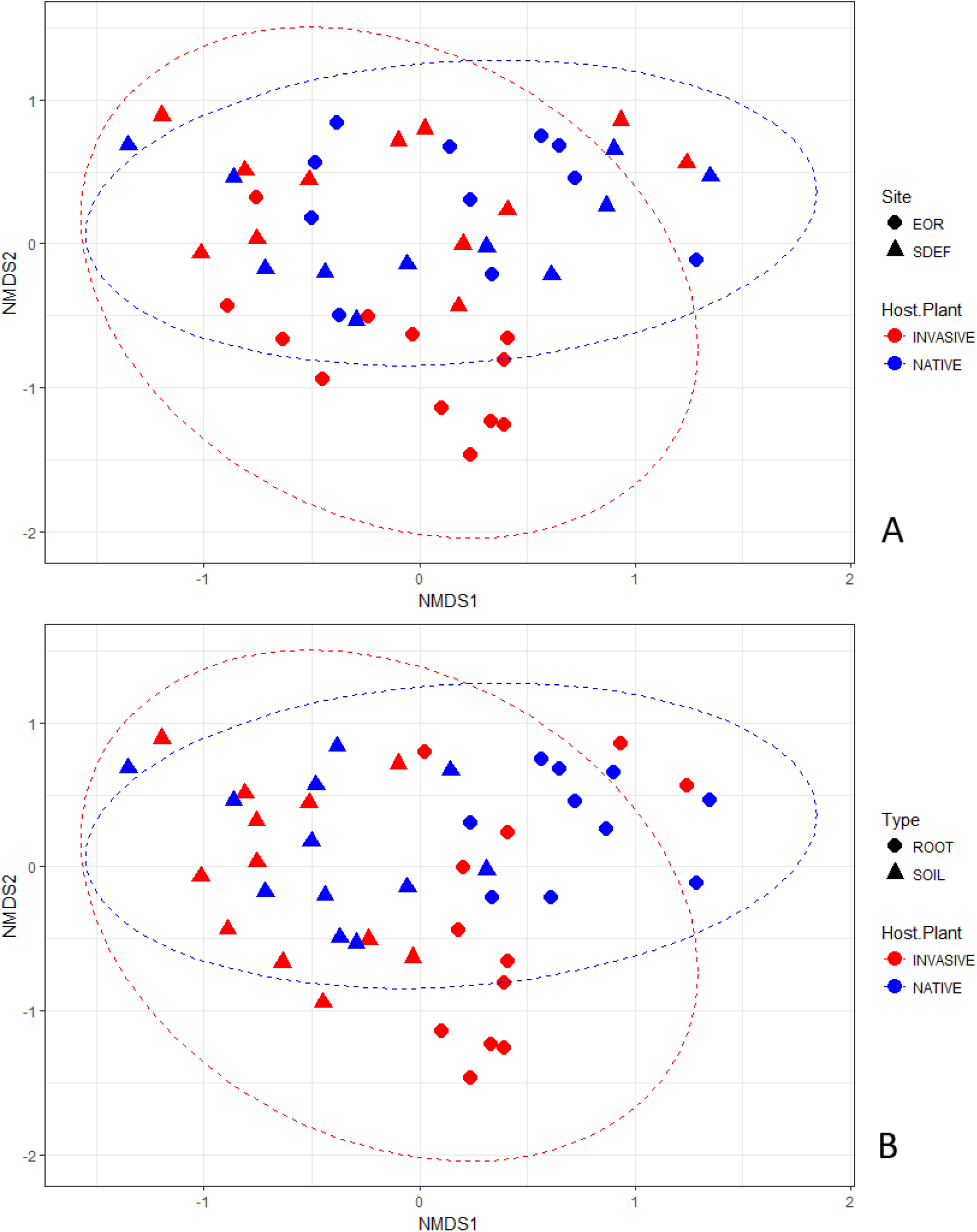
AMF (SSU) Bray-Curtis NMDS plots. In panel A, color is host plant and shape denotes site: San Dimas Experimental Forest (SDEF) or Emerson Oaks Reserve (EOR). In panel B color denotes host plant and shape denotes if the community is from a root or soil sample.

**Figure 2:**
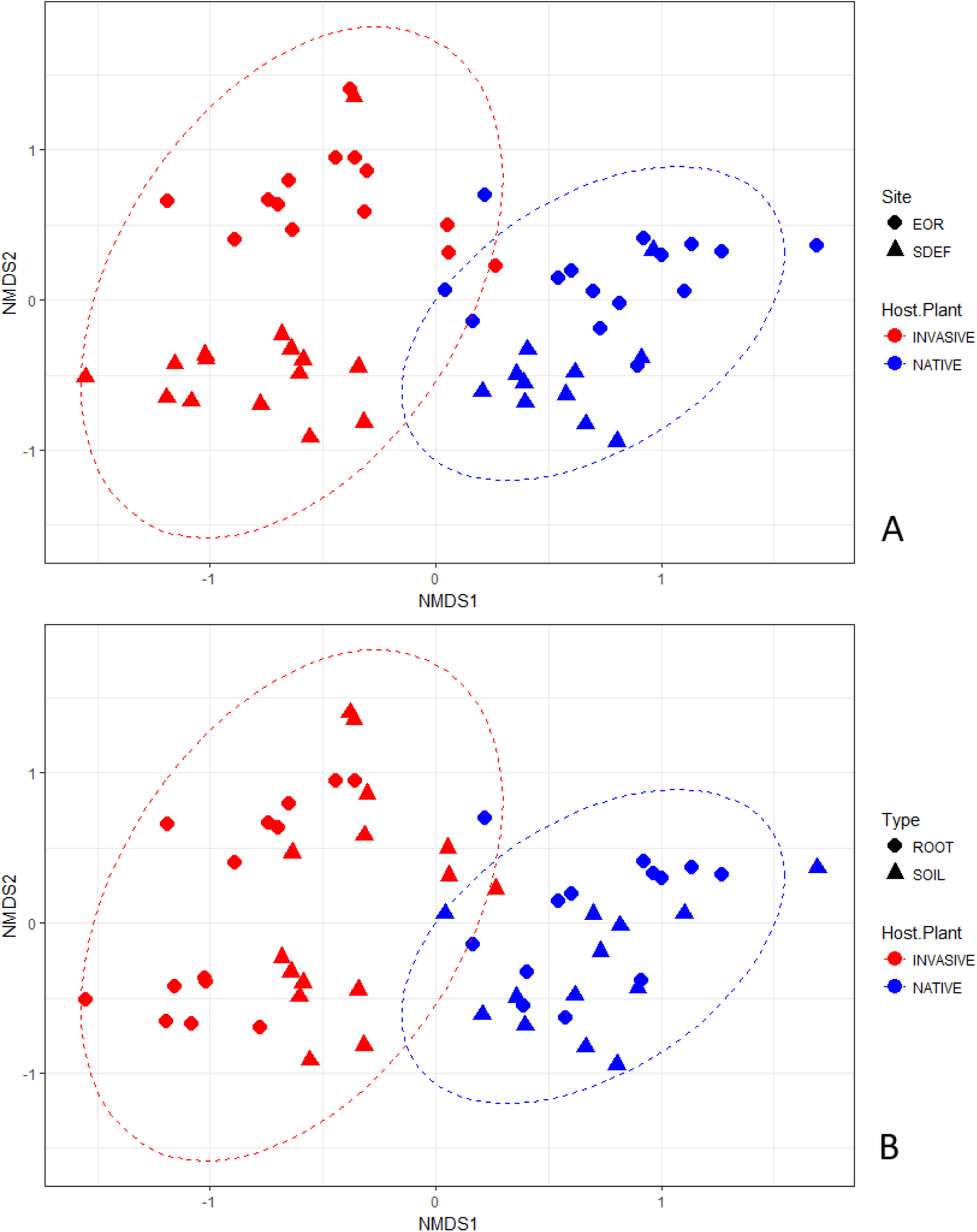
General Fungal Community (ITS2) Bray-Curtis NMDS plots. In panel A, color is host plant and shape denotes site: San Dimas Experimental Forest (SDEF) or Emerson Oaks Reserve (EOR). In panel B color denotes host plant and shape denotes if the community is from a root or soil sample.

### Functional group responses

#### Rhizophilic AMF

Richness and relative read abundance of rhizophilic AMF was greater in native than invasive roots (P = 0.008 and 0.02, respectively; Figure 3a; Table S1). Rhizophilic AMF richness and relative abundance in roots was negatively correlated with soil NH_4_ concentrations (P = 0.003 and 0.016, respectively; Table S1). Rhizophilic AMF richness and relative read abundance in roots were positively associated with soil NO_3_ concentrations (P = 0.01 and 0.002, respectively; Table S1). There were no differences in the richness or relative abundance of rhizophilic taxa in soils underneath native shrubs and invasive grasses (P = 0.71 and 0.77, respectively; Figure 3b).

**Figure 3:**
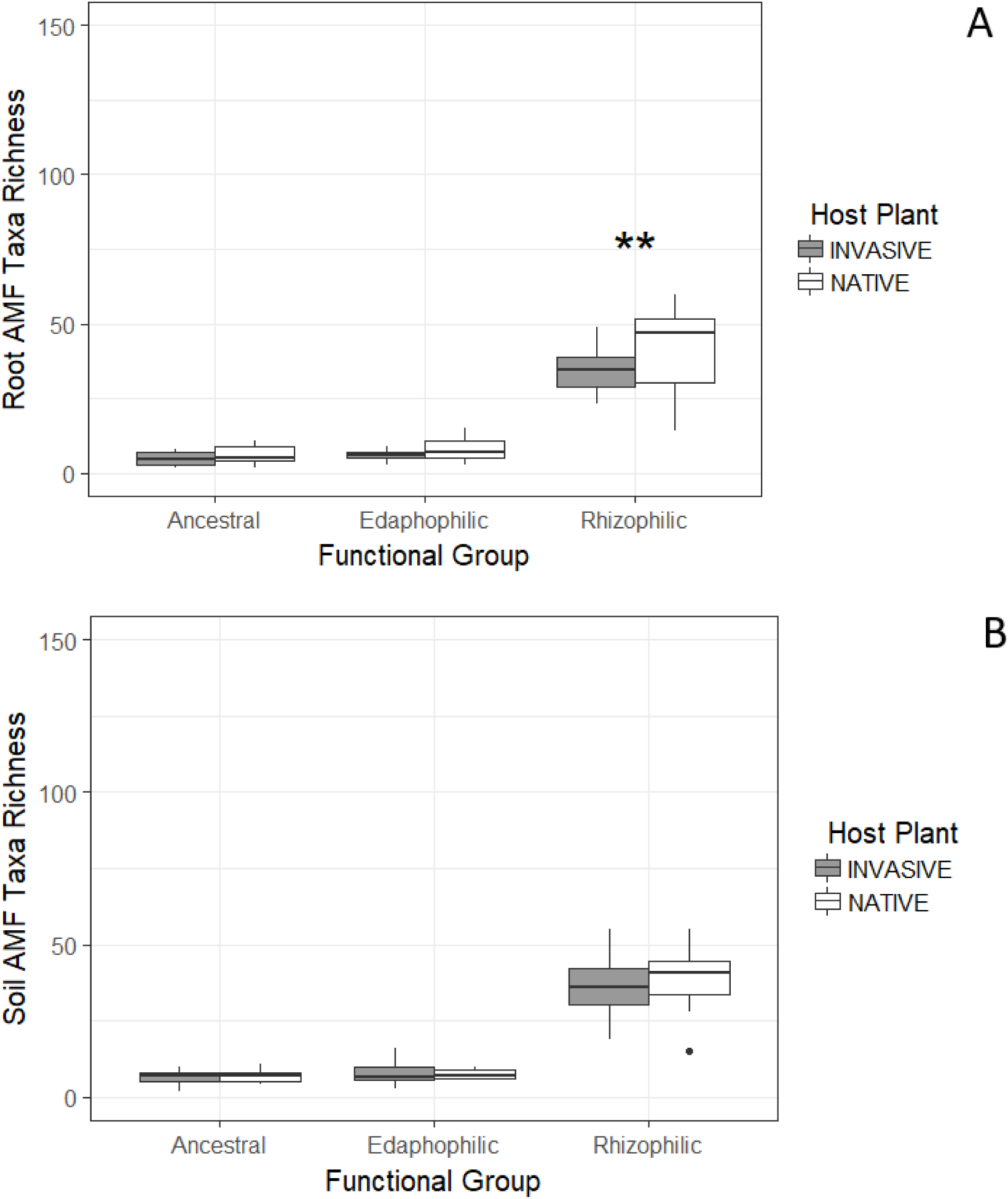
SSU or arbuscular mycorrhizal fungal (AMF) root (A) and soil (B) communities by functional group by aggregating species by family using the phylogenetic scheme in Table 1. AMF taxa richness is the number of times a unique taxonomic unit is encountered in each sample. *** denotes significant difference by host plant type at P < 0.001, ** denotes significance at P < 0.01 and * denotes significance at P < 0.05 from GLM outputs in table S1.

#### Edaphophilic AMF

The relative abundance of edaphophilic AMF was higher in native shrub roots than in invasive grass roots (P = 0.02; Figure 3a; Table S1), while richness did not differ between these plant roots (P = 0.26; Figure 3b). The richness of edaphophilic AMF in soils underneath native shrubs and invasive grasses did not differ (P = 0.77), however edaphophilic AMF were relatively more abundant in native soils (P = 0.007, Table S1). Richness of edaphophilic AMF in roots was positively correlated with soil NO_3_ (P = 0.04; Table S1). Relative abundance of edaphophilic AMF in soils was negatively correlated with soil NH_4_ concentrations and positively correlated with soil NO_3_ concentrations (P = 0.03 and 0.005, respectively; Table S1).

#### Ancestral AMF

Native roots had greater relative read abundance, but not richness of ancestral AMF families when compared to invasive (P = 0.006 and 0.2, respectively; Table S1). Host plant was not included in the ancestral soil relative abundance and richness models after model selection. Root ancestral AMF richness was negatively correlated with soil NH_4_ concentrations and positively associated with soil NO_3_ concentrations (P = 0.01 and 0.01; Table S1). Conversely, soil ancestral AMF richness and relative read abundance were negatively associated with increased soil NO_3_ concentrations (P = 0.003 and 0.03, respectively; Table S1).

#### Non-AMF Symbionts

Non-AMF symbionts – including EMF – had greater richness (Figure 4a) and relative abundance in native roots (P = 0.002 and 0.003; Table S1). Non-AMF symbiont richness, but not abundance, was also greater in native soils (Figure 4b, P = 0.035 and 013, respectively; Table S1). Non-AMF symbiont richness in roots was negatively associated with soil NH_4_ and NO_3_ concentrations (P = 0.001 and 0.001, respectively; Table S1). Conversely, non-AMF symbiont relative abundance was positively associated with soil NH_4_ and NO_3_ soil concentration (P = 0.001 and 0.003, respectively; Table S1).

**Figure 4:**
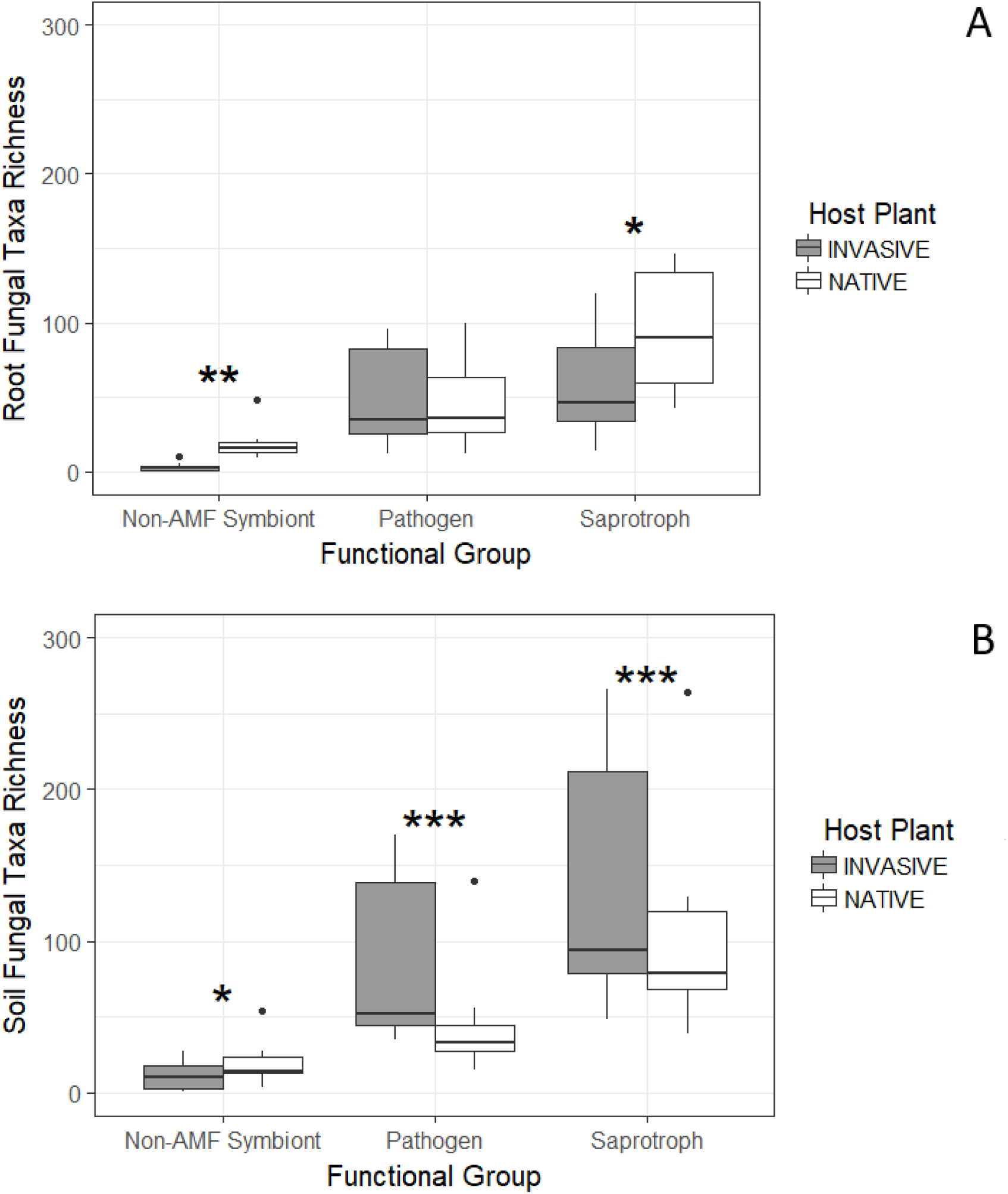
ITS or general fungal community root (A) and soil (B) communities by functional group by aggregating species using FUNguild. Fungal taxa richness is the number of times a unique taxonomic unit is encountered in each sample. *** denotes significant difference by host plant at P < 0.001, ** denotes significance at P < 0.01 and * denotes significance at P < 0.05 from GLM outputs in table S1.

#### Pathogens

Pathogen fungi were relatively more abundant in invasive grass roots (Figure 4a, P = 0.011; Table S1), however richness did not differ (Figure 4b, P = 0.63). Pathogen richness (Figure 4b) and relative abundance were greater in invasive soils (P = 0.001 and 0.001, respectively; Table S1). SDEF had higher pathogen richness and relative abundance in soils than EOR (P = 0.001 and 0.001; Table S1). SDEF had higher pathogen richness and relative abundance in soils than EOR (P = 0.001 and 0.001; Table S1).

#### Saprotrophs

Saprotroph relative abundance was greater in invasive soils (P = 0.001), however saprotroph richness was greater in native soils (P = 0.001, Figure 4b; Table S1). Richness and relative abundance of saprotrophs in soils were positively associated with higher soil NH_4_ concentration (P = 0.001, 0.001; Table S1). Saprotroph richness in soils negatively correlated with soil NO_3_ concentration (P = 0.022; Table S1). Root saprotroph richness was higher in native roots when compared to invasive (P = 0.03; Table S1).

### Taxonomic abundance responses

#### AMF Families

The relative abundance of AMF families did not vary significantly between the roots nor soils beneath invasive grasses and native shrubs (Tables S4 and S5). Taxa belonging to: Archaeosporaceae, Claroideoglomeraceae, Diversisporaceae, and Glomeraceae were relatively more abundant in roots at EOR (P ≤ 0.1, Table S4), however we found no significant differences between sites in soils (Table S5). Relative read abundance for all AMF families in roots was positively correlated with soil NO_3_ concentrations (P ≤ 0.1, Table S4). We observed increases in relative abundance of Acaulosporaceae, Archaeosporaceae, Claroideoglomeraceae, Diversisporaceae, Glomeraceae, and Paraglomeraceae in roots with increasing soil P concentrations (P ≤ 0.1, Table S4). In soils, fewer environmental variables were significantly associated with relative abundance of AMF families. Relative abundance of taxa belonging to: Acaulosporaceae, Archaeosporaceae, Diversisporaceae, and Paraglomeraceae were positively associated with soil pH concentrations ranging from 6 to 7 (P ≤ 0.1, Table S5). Relative abundance of Acaulosporaceae, Ambisporaceae, and Claroideoglomeraceae in soils increased with increasing soil NH_4_ concentrations (P ≤ 0.1, Table S5).

#### Variance partitioning

Environmental predictors (host plant, site, NH_4_, NO_3_, pH, and P) explained 92%±7% of the variance in the AMF root community model (Figure 5a, Table S6). Relative abundance of Ambisporaceae in roots, which was more abundant in native samples, had the most model variance explained by host plant, 19%, and for all other AMF families host plant explained less than 10% of model variance (Table S6, Figure 5a). Soil NO_3_ concentrations explained the largest amount of model variance in the root model (33%± 4%, Figure 5a, Table S6). In soil communities, total environmental predictors explained 92%±7% of model variance (Figure 5b, Table S7). Soil P concentrations explained the largest amount of the variance ranging from 35% ± 14% of the variation in the soil model (Figure 5b, Table S7).

**Figure 5:**
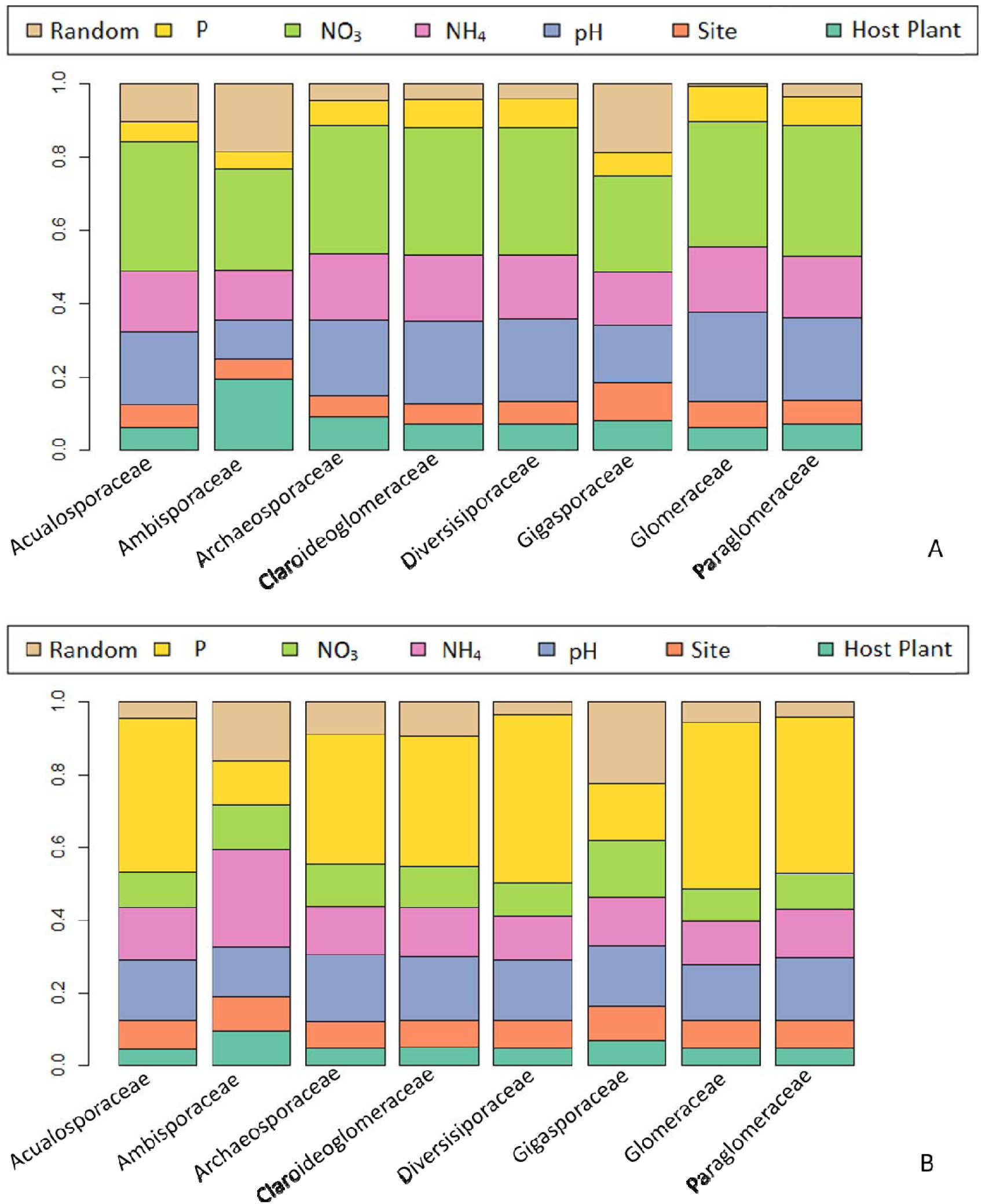
Variance partitioning of AMF (SSU) relative of abundance in the HMSC models of root (A) and soil (B) communities at the family level. Variation is partitioned by fixed effects (P, NO_3_, NH_4_, pH, site, and host plant) as well as the random effect (individual sampled).

## Discussion

The increased proportion of edaphophilic AMF among native shrub roots and soils provides support for our first hypothesis, and is consistent with other studies in which locally adapted fungi exhibit a preference for locally adapted host plants (Johnson *et al.* 2009). We expected that invasive grasses would host more rhizophilic AMF taxa, however these taxa were more abundant and rich in native shrub roots. We hypothesized that invasive grasses would harbor fewer pathogens but did not find strong support for this. Instead, we found that pathogenic fungi were relatively more abundant in invasive roots and soils. Microscopic observations showed that invasive grass roots were colonized by both AMF and non-AMF at higher rates than the roots of the native shrub *Adenostoma fasciculatum*. We expected that invasive hosts would interact with soil N, resulting in decreased richness and abundance of edaphophilic AMF, but we found little support for this hypothesis. Relative abundance of edaphophilic AMF in soils was negatively correlated with soil NH_4_ concentrations, but positively correlated with soil NO_3_ concentrations. Richness of rhizophilic, edaphophilic, and ancestral AMF in roots were positively correlated with soil NO_3_ concentrations. Overall these findings suggest that while the same pool of AMF mutualists is available for both *A. fasciculatum* and invasive grasses, the mycorrhizas formed between these plants and AM fungi differ, potentially because of differences in plant roots and fungal biomass allocation (Maherali & Klironomos 2007; Powell *et al.* 2009; Sikes *et al.* 2009, 2010).

### Functional group responses

#### Symbiotic fungi

Lower richness and relative abundance of some AMF functional groups in invasive roots, concurs with past research suggesting that invasive annual grasses may be less dependent on AMF mutualisms (Allen 1984; Richardson *et al.* 2000; Callaway *et al.* 2004; Reinhart & Callaway 2006; Busby *et al.* 2011, 2013). If invasive grasses are less dependent on soil mutualists, this could facilitate rapid establishment of these grasses following disturbance. The degraded mutualist hypothesis purports that invasive plant species that successfully establish due to decreased dependence on soil mutualisms will decrease the presence of plant species that are highly dependent on mutualisms over time (Vogelsang & Bever 2009). We found relative decreases in three groups of soil symbionts associated with invasion: non-AMF symbionts (including EMF), edaphophilic and rhizophilic AMF. This suggests that invasive persistence may decrease the presence of multiple soil symbionts that native species depend on for pathogen protection and for increased access to soil resources. However, we also observed higher rates of AMF colonization in invasive than native roots. Invasive samples had more OTUs that could not be assigned taxonomy in MaarjAM (‘no blast hit’) than native samples, which could mean that there were shifts in the abundance of AMF taxa that may not have been observed in our sequence data. Another possibility is that the ‘no blast hit’ may be a eukaryote other than AMF, because QIIME requires a 90% similarity to assign a match and AMF are relatively conserved (Powell *et al.* 2009).

We observed decreases in relative abundance coupled with decreases in richness for some groups of AMF, which may result in losses of necessary function and/or taxa native plants rely on. Specifically, decreases in proportions of edaphophilic AMF would decrease the presence of extraradical hyphae that *A. fasciculatum* depends on for resource uptake. These results, combined with no change in richness associated with invasion, align with previous findings in the literature that variation in AMF composition between systems is often due to differences in abundance rather than a distinct taxonomic composition (Hart *et al*. 2016; Hijri *et al.* 2006; Öpik *et al.* 2008). This suggests that under invasion we may see shifts in the relative abundance of taxa, but not a complete turnover of AMF taxa that are present.

Our results suggest that differences in richness and relative abundance of symbionts, both AMF and non-AMF, may be associated with host plant identity. Non-AMF symbionts detected by ITS2 sequencing were mainly EMF indicating their presence even though they were not detected microscopically. Nevertheless, *A. fasciculatum* forms EMF under wet conditions (Allen *et al.* 1999), and invasive grass encroachment may indirectly decrease EMF colonization by rapidly depleting soil moisture (Melgoza *et al.* 1990). It may be important to understand the richness and abundance of different functional groups of fungi in natural recolonization or restoration efforts of slow-growing shrubs like *A. fasciculatum*, that could be highly dependent on locally diverse adapted symbiotic relations for establishment (Azcón-Aguilar *et al*., 2003; Johnson *et. al.,* 2009).

#### Pathogenic fungi and other non-AMF fungi

We did not find strong evidence to support pathogen release in this system (Mitchell & Power 2003; Kardol *et al.* 2007; Van Grunsven *et al.* 2007; Reinhart *et al.* 2010), as pathogen relative abundance was greater in invasive roots and soils. SDEF had a greater richness of pathogens when compared to EOR, which may be due to increased soil N availability at SDEF. Additionally, we observed greater relative abundance of rhizophilic AMF in soils and richness in roots at SDEF which may promote greater pathogen protection (Maherali & Klironomos 2007; Sikes *et al.* 2009). It is important to note that in using FUNguild to assign functional groups while also filtering out all taxa with the confidence level ‘possible’ (Nguyen *et al.* 2016), we lost potentially valuable data. However, using only conservative functional group assignments with the confidence levels ‘highly probable’ and ‘probable’ protected the integrity of our interpretations.

There was an increase in non-AMF colonization in invasive roots that could be due to increased pathogen or saprotrophic colonization. This was also supported by ITS2 data, which showed significant differences in pathogen and saprotrophic richness or relative abundance in invasive grass roots. However, we observed greater AMF colonization in invasive than native roots which may confer greater pathogen protection (Maherali & Klironomos 2007; Sikes *et al.* 2009). Coarse roots of *A. fasciculatum* were predominant in our samples, which may have contributed to our observations of lower overall colonization in shrub than fine grass roots. Another study reported higher rates of AMF colonization in *A. fasciculatum* as well as EMF in wet but not dry years (Allen *et al.* 1999). We sampled during a drought year which likely decreased the presence of EMF in these soils.

Recent research suggests that some fungi may have the potential to occupy complex or multiple niches (Glynou *et al.* 2017; Selosse *et al.* 2018). Our findings of greater saprotroph richness in *A. fasciculatum* living roots support this by indicating that some fungi could be acting as opportunistic pathogens. The idea that fungi possess dual niches stems from the evolutionary propensity of fungi to shift ecological niches, while often retaining their previous niche (Selosse *et al.* 2018). Therefore, these presumably saprotrophic fungi may be acting as facultative pathogens in roots and saprotrophs in soils. Additionally, invasive annual grasses produce larger amounts of easily decomposed litter, which agrees with our observations of greater relative abundance of saprotrophs in invasive associated soils (De Deyn *et al.* 2008).

We used a recently developed AMF guild framework and FUNGuild to assign function to fungal taxa that we observed aimed at understanding the ecological relevance of taxonomic differences between host plants and across environmental conditions. Out of necessity for interpretation, both methods constrain descriptions of fungal function to simple categories. Despite this need, it is important to remember that interactions between fungi and plant hosts are complex, varying within taxa and individuals, with the potential to occupy multiple ecological niches under varying environmental conditions (Selosse *et al.* 2018). Thus, both the AMF guild framework and the FUNGuild application that we use in this study are coarse tools which at best approximate fungal ecological functioning. Our approach is supported by Treseder et al. (2018), who found that high soil N was negatively related to external hyphal length. The use of sequencing data to understand fungal ecology is ultimately limited by research that links fungal life histories and ecological functioning to sequence data.

### Taxonomic responses

#### AMF families

We did not observe effects of site or host plant on any AMF families in roots or soils, but in our functional guild analyses we found that rhizophilic and edaphophilic AMF were relatively more abundant in native roots. This indicates that the complexity of family-level community composition may be effectively reduced using a functional grouping approach, allowing nuanced relationships between invasion and AMF communities to be resolved at this scale. However, variance partitioning during family-level analysis indicated that environmental variables differentially structure AMF root and soil communities. For soils, the largest amount of variability across all AMF families was attributed to soil P concentrations. However, less variability was explained for Gigasporaceae and Ambisporaceae abundance by soil P compared to other AMF families. The Gigaposraceae family falls into the edaphophilic AMF group, but the Diversisporaceae, the other family in this group, has much more variability explained by soil P. This may mean that responses to environmental variables are not consistent across resource allocation strategies of AMF, or that we still need a better understanding of resource allocation of some families.

For roots, the largest amount of variability across all AMF families was attributed to soil NO_3_ concentrations, meaning that selectivity of the host plant and fungi in initializing mutualisms may heavily depend on this. We observed increases in abundance for most AMF families with increased soil NO_3_. Specifically, Glomeraceae and Paraglomeraceae appear to be the most positively associated with the higher soil NO_3_ concentrations, whereas Gigasporaceae and Ambisporaceae showed little increase with elevated NO_3_, a pattern that was also observed by Egerton-Warburton and Allen (2000) and Treseder et al. (2018). Interestingly, Glomeraceae and Paraglomeraceae fall into the rhizophilic AMF group while Gigasporaceae is edaphophilic. This suggests that edaphophilic AMF that mainly produce extraradical hyphae are more sensitive to increasing soil N concentrations, whereas rhizophilic AMF that primarily produce intraradical hyphae may be stimulated by increased soil N.

## Conclusions

Invasion decreased the abundance and richness of both AMF and non-AMF symbionts, suggesting that type conversion from native shrubland to non-native grasses may decrease the richness and abundance of some symbiotic fungal taxa in soils (Busby *et al.*, 2011; Busby *et al.*, 2013; Hawkes *et al.*, 2006). We observed differences in relative abundance and richness of functional groups of AMF between native and invasive root and soil communities. However, in our taxonomic analyses we did not find differences in abundance of any AMF family between native and invasive roots or soils. Our results support the hypothesis that native shrubs host a more abundant (but not richer) community of edaphophilic AMF. Decreases in available edaphophilic AMF taxa may hamper the re-establishment of native shrubs into their home range by decreasing access to host-specific mutualists (Johnson *et al.* 2009). Our results do not support our hypothesis that invasive grasses would host more rhizophilic taxa, as rhizophilic AMF were 14 richer and relatively more abundant in native shrub roots. However, we did observe a larger amount of both AMF and non-AMF colonization in invasive grass roots.

Previous work on soil fungal communities and invasion provides substantive evidence for pathogen release in other systems (Mitchell & Power 2003; Kardol *et al.* 2007; Van Grunsven *et al.* 2007; Reinhart *et al.* 2010). Our hypothesis that pathogen release is promoting high abundances of invasive plants in chaparral is contradicted by higher relative abundances of pathogens in invasive plant roots, coupled with higher rates of non-mycorrhizal root colonization. The higher relative abundances of these potentially parasitic fungi in invasive grass roots compared to native shrubs may be a result of density dependence, given that invasive grasses occur at higher densities than native shrubs. Future work would benefit from (i) confirming that these potential parasites negatively affect invasive plants and (ii) investigating invasive plant and parasitic fungal abundance dynamics over multiple seasons.

We did not find strong support for our hypothesis that elevated soil N concentrations would reduce the relative abundance of edaphophilic AMF. Surprisingly, we observed both positive and negative interactions between increased soil N and edaphophilic AMF richness and relative abundance. However, we did observe decreased relative abundance of edaphophilic AMF associated with invasive hosts. Future work should include experimental manipulation of soil N and invasion to better resolve the relationship between N availability, exotic plant invasion, and AMF composition. Our results illustrate the importance of including both microscopic observations and sequencing data in efforts to understand AMF. There is a need for more information about the relationship between taxonomy and function of both AMF and other fungi, to address how the interplay of fungi and plants will shift in response to global change.

## Acknowledgements

We thank Jeffrey Diez and Courtney Collins, University of California Riverside, for guidance on statistical analyses as well as Catherine Gehring, Northern Arizona University, for facilitating laboratory work. We thank Tye Morgan at USDA-ARS Soils Laboratory, Reno, NV for assistance with processing soils for bicarbonate-extractable P. We thank members of the Allen and Aronson labs for comments and feedback on the manuscript. We also would like to thank two anonymous reviewers for their feedback, which helped to improve this manuscript. Funding was provided by University of California Riverside, Department of Botany and Plant Sciences and the Center for Conservation Biology, as well as a graduate fellowship from NASA MUREP Institutional Research Opportunity (MIRO) grant number NNX15AP99A, and a graduate fellowship from the University of California’s Institute for the Study of Ecological and Evolutionary Climate Impacts (ISEECI), funded by a UC Presidential Research Catalyst Award.

## References

Allen, E.B. & Allen, M.F. (1980) Natural Re-Establishment of Vesicular-Arbuscular Mycorrhizae Following Stripmine Reclamation in Wyoming. Journal of Applied Ecology, 17, 139–147.

Allen, E.B. 1984. Mycorrhizae and colonizing annuals: implications for growth, competition, and succession. S. E. Williams and M. F. Allen, eds. VA Mycorrhizae and Reclamation of Arid and Semiarid Lands. University of Wyoming Agr. Expt. Sta. Laramie, Wyoming, 42–52.

Allen, E.B., Allen, M.F.A., Corkidi, L., Egerton-warburton, L. & Gomez Pompa, A. (2003) Impacts of Early- and Late-Seral Mycorrhizae During Restoration in Seasonal Tropical Forest, Mexico. Ecological Applications, 13, 1701–1717.

Allen, M.F., Egerton-Warburton, L.M., Allen, E.B. & Karen, O. (1999) Mycorrhizae in Adenostoma fasciculatum Hook. and Arn.: A combination of unusual ecto- and endo-forms. Mycorrhiza, 8, 225–228.

Altschul, S.F., Gish, W., Miller, W., Myers, E.W. & Lipman, D.J. (1990) Blast Local Alignment Search Tool. Journal of Molecular Biology, 403–410.

Alvarado, P., Teixeira, M., Andrews, L., Fernandez, A., Santander, G., Doyle, A.L., Perez, M., Yegres, F., Mendoza, M. & Barker, B.M. (2017) Molecular detection of Coccidioides posadasii and associated fungal communities from soils in endemic areas of coccidioidomycosis in Venezuela. In Press.

Amend, A.S., Martiny, A.C., Allison, S.D., Berlemont, R., Goulden, M.L., Lu, Y., Treseder, K.K., Weihe, C. & Martiny, J.B.H. (2015) Microbial response to simulated global change is phylogenetically conserved and linked with functional potential. ISME J, 10, 109–118.

Andrews S. (2010) FastQC: a quality control tool for high throughput sequence data. Available online at: http://www.bioinformatics.babraham.ac.uk/projects/fastqc||

Aronesty, E. (2011) eautils: “Command-line tools for processing biological sequencing data.”

Ashbacher, A. & Cleland, E. (2015) Native and exotic plant species show differential growth but similar functional trait responses to experimental rainfall. Ecosphere, 6, 1–14.

Azcón-Aguilar, C., Palenzuela, J. Roldán, Bautista, S., Vallejo, R., Barea, J.M., (2003) Analysis of the mycorrhizal potential in the rhizosphere of representative plant species from desertification-threatened Mediterranean shrublands. Applied Soil Ecology, 22(1), 29–37.

Bengtsson-Palme, J., Ryberg, M., Hartmann, M., Branco, S., Wang, Z., Godhe, A., Wit, P. De, Marisol, S., Ebersberger, I., Sousa, F. De, Amend, A.S., Jumpponen, A., Unterseher, M., Kristiansson, E., Abarenkov, K., Bertrand, Y.J.K., Sanli, K., Eriksson, K.M., Vik, U., Veldre, V. & Nilsson, R.H. (2013) Improved software detection and extraction of ITS1 and ITS2 from ribosomal ITS sequences of fungi and other eukaryotes for analysis of environmental sequencing data. Methods in Ecology and Evolution, 4, 914–919.

Berry, D., K. Ben Mahfoudh, M. Wagner, and A. Loy (2012). Barcoded Primers Used in Multiplex Amplicon Pyrosequencing Bias Amplification. Applied and Environmental Microbiology 78, 612.

Blanchet, F.G. & Tikhonov, G. (2016). HMSC: Hierarchical Modelling of Species Community. R package version 2.0-0.

Busby, R.R., Gebhart, D.L., Stromberger, M.E., Meiman, P.J. & Paschke, M.W. (2011) Early seral plant species’ interactions with an arbuscular mycorrhizal fungi community are highly variable. Applied Soil Ecology, 48, 257–262.

Busby, R.R., Stromberger, M.E., Rodriguez, G., Gebhart, D.L. & Paschke, M.W. (2013) Arbuscular mycorrhizal fungal community differs between a coexisting native shrub and introduced annual grass. Mycorrhiza, 23, 129–41.

Callaway, R. M., D. Cipollini, K. Barto, G. C. Thelen, S. G. Hallett, D. Prati, K. Stinson, *and* J. Klironomos. (2008) Novel weapons: invasive plant suppresses fungal mutualists in America but not in its native Europe. Ecology, 89, 1043–1055.

Callaway, R.M., Thelen, G.C., Barth, S., Ramsey, P.W. & Gannon, J.E. (2004) SOIL FUNGI ALTER INTERACTIONS BETWEEN THE INVADER CENTAUREA MACULOSA AND NORTH AMERICAN NATIVES. Ecology, 85, 1062–1071.

Caporaso, J.G., Kuczynski, J., Stombaugh, J., Bittinger, K., Bushman, F.D., Costello, E.K., Fierer, N., Peña, A.G., Goodrich, J.K., Gordon, J.I., Huttley, G. a, Kelley, S.T., Knights, D., Koenig, J.E., Ley, R.E., Lozupone, C. a, Mcdonald, D., Muegge, B.D., Pirrung, M., Reeder, J., Sevinsky, J.R., Turnbaugh, P.J., Walters, W. a, Widmann, J., Yatsunenko, T., Zaneveld, J. & Knight, R. (2010) correspondence QIIME allows analysis of high-throughput community sequencing data Intensity normalization improves color calling in SOLiD sequencing. Nature Publishing Group, 7, 335–336.

D’Antonio, C.M. & Vitousek, P.M. (1992) Biological Invasions by Exotic Grasses, the Grass/Fire Cycle, and Global Change. Annual Review of Ecology and Systematics, 23, 63–87.

De Deyn, G.B., Cornelissen, J.H.C. & Bardgett, R.D. (2008) Plant functional traits and soil carbon sequestration in contrasting biomes. Ecology letters, 11, 516–31.

Dumbrell, A. J., P. D. Ashton, N. Aziz, G. Feng, M. Nelson, C. Dytham, A. H. Fitter, and T. Helgason. (2011) Distinct seasonal assemblages of arbuscular mycorrhizal fungi revealed by massively parallel pyrosequencing-Supplement. The New Phytologist 190, 794–804.

Dunn, P.H., Barro, S.C., Wells, W.G., Poth, M.A., Wohlgemuth, P.M., Colver, C.G., 1988. The San Dimas Experimental Forest□: 50 Yearsof Research, Forest Service: General Technical Report.

Egerton-Warburton, L.M. & Allen, E.B. (2000) SHIFTS IN ARBUSCULAR MYCORRHIZAL COMMUNITIES ALONG AN ANTHROPOGENIC NITROGEN DEPOSITION GRADIENT. Ecological Applications, 10, 484–496.

Egerton-Warburton, L.M., Graham, R.C., Allen, E.B. & Allen, M.F. (2001) Reconstruction of the historical changes in mycorrhizal fungal communities under anthropogenic nitrogen deposition. Proceedings. Biological sciences / The Royal Society, 268, 2479–2484.

Egerton-Warburton, L.M., Johnson, N.C. & Allen, E.B. (2007) MYCORRHIZAL COMMUNITY DYNAMICS FOLLOWING NITROGEN FERTILIZATION□: A CROSS-SITE TEST IN FIVE GRASSLANDS. Ecological Monographs, 77, 527–544.

Fenn, M.E., Allen, E.B., Weiss, S.B., Jovan, S., Geiser, L.H., Tonnesen, G.S., Johnson, R.F., Rao, L.E., Gimeno, B.S., Yuan, F., Meixner, T. & Bytnerowicz, A. (2010) Nitrogen critical loads and management alternatives for N-impacted ecosystems in California. Journal of Environmental Management, 91, 2404–2423.

Glynou, K., Nam, B., Thines, M. & Maci, J.G. (2017) Facultative root-colonizing fungi dominate endophytic assemblages in roots of nonmycorrhizal *Microthlaspi* species. New Phytologist, 217, 1190–1202.

Van Grunsven, R.H.A., van der Putten, W.H., Bezemer, T.M., Tamis, W.L.M., Berendse, F. & Veenendaal, E.M. (2007) Reduced plant-soil feedback of plant species expanding their range as compared to natives. Journal of Ecology, 95, 1050–1057.

Hart, M.M. & Reader, R.J. (2002) Taxonomic basis for variation in the colonization strategy of arbuscular mycorrhizal fungi. New Phytologist, 153, 335–344.

Hart, M.M., Zaitsoff, P.D., van der Heyde, Pither, J. (2016) Testing life history and trait-based predictions of AM fungal community assembly. Pedobiologia-J. Soil Ecol.

Hausmann, N.T. & Hawkes, C. V. (2009) Plant neighborhood control of arbuscular mycorrhizal community composition. New Phytologist, 183, 1188–1200.

Hawkes, C. V., Belnap, J., D’Antonio, C., Firestone, M.K., (2006). Arbuscular mycorrhizal assemblages in native plant roots change in the presence of invasive exotic grasses. Plant and Soil, 281(1-2), 69–380.

Hijri, I., Sýkorová, Z., Oehl, F., Ineichen, K., Mäder, P., Wiemken, A., Redecker, D., (2006) Communities of arbuscular mycorrhizal fungi in arable soils are not necessarily low. Molecular Ecology, 15, 2277–2289

Hooper, D.U. & Vitousek, P.M. (1998) Effects of Plant Composition and Diversity on Nutrient Cycling. Ecological Monographs, 68, 121–149.

Inderjit & van der Putten, W.H. (2010) Impacts of soil microbial communities on exotic plant invasions. Trends in Ecology and Evolution, 25, 512–519.

Johnson, N.C., Wilson, G.W.T., Bowker, M.A., Wilson, J.A. & Miller, R.M. (2009) Resource limitation is a driver of local adaptation in mycorrhizal symbioses. Proceedings of the National Academy of Sciences of the United States of America, 107, 2093–2098.

Kardol, P., Cornips, N.J., van Kempen, M.M.L., Bakx-Schotman, J.M.T. & van der Putten, W.H. (2007) Microbe-mediated plant–soil feedback causes historical contingency effects in plant community assembly. Ecological Monographs, 77(2), 147–162.

Kõljalg, U., Nilsson, R.H., Abarenkov, K., Tedersoo, L., Taylor, A.F.S., Bahram, M., Bates, S.T., Bruns, T.D., Bengtsson-Palme, J., Callaghan, T.M., Douglas, B., Drenkhan, T., Eberhardt, U., Dueñas, M., Grebenc, T., Griffith, G.W., Hartmann, M., Kirk, P.M., Kohout, P., Larsson, E., Lindahl, B.D., Lücking, R., Martín, M.P., Matheny, P.B., Nguyen, N.H., Niskanen, T., Oja, J., Peay, K.G., Peintner, U., Peterson, M., Põldmaa, K., Saag, L., Saar, I., Schüßler, A., Scott, J. a., Senés, C., Smith, M.E., Suija, A., Taylor, D.L., Telleria, M.T., Weiss, M. & Larsson, K.H. (2013) Towards a unified paradigm for sequence-based identification of fungi. Molecular Ecology, 22, 5271–5277.

Kormanik, P. P. & McGraw, A. C. (1982). Quantification of vesicular-arbuscular mycorrhizas in plant roots. In Methods and Principles of Mycorrhizal Research (Ed. by N. C. Schenck), The American Phytopathological Society, St Paul, Minnesota, 37–45.

Koske, R.E. & Gemma, J.N. (1989) SHORT COMMUNICATIONS A modified procedure for staining roots to detect VA mycorrhizas. Mycological Research, 92, 486–488.

Lee, J., S. Lee, and J. P. W. Young. 2008. Improved PCR primers for the detection and identification of arbuscular mycorrhizal fungi. FEMS Microbiology Ecology, 65, 339–349.

Mahe, F., Rognes, T., Quince, C., Vargas, C. De & Dunthorn, M. (2014) Swarm□: robust and fast clustering method for amplicon-based studies. PeerJ, 1–13.

Maherali, H. & Klironomos, J.N. (2007) Influence of phylogeny on fungal community assembly and ecosystem functioning. Science, 316, 1746–8.

Martin, M. (2011) Cutadapt Removes Adapter Sequences from High-Throughput Sequencing Reads. EMBnet.journal, 17, 5–7.

McGonigle, T.P., Miller, M.H., Evans, D.G., Fairchild, G.L. Swan, J.A.,(1990) A new method which gives an objective measure of colonization of roots by vesicular-arbuscular mycorrhizal fungi. New Phytologist, 115(3), 495–501.

Melgoza, G., Nowak, R.S. & Tausch, R.J. (1990) Soil water exploitation after fire□: competition between Bromus tectorum (cheatgrass) and two native species. Oecologia, 83, 7–13.

Mitchell, C.E. & Power, A.G. (2003) Release of invasive plants from fungal and viral pathogens. Nature, 421, 625–7.

Müller, G., Horstmeyer, L., Rönneburg, T., van Kleunen, M. & Dawson, W. (2016) Alien and native plant establishment in grassland communities is more strongly affected by disturbance than above- and below-ground enemies. Journal of Ecology, 104, 1233–1242.

Nguyen, N.H., Song, Z., Bates, S.T., Branco, S., Tedersoo, L., Menke, J., Schilling, J.S. & Kennedy, P.G. (2016) FUNGuild□: An open annotation tool for parsing fungal community datasets by ecological guild. Fungal Ecology, 20, 241–248.

Nilsson, R.H., Tedersoo, L., Ryberg, M., Kristiansson, E., Hartmann, M., Unterserher, M., Porter, T., Bengtsson-Palme, J., Walker, D., De Sousa, F., Gamper, H.A., Larsson, E., Larsson, K.H., Koljalg, U., Edgar, R.C. & Abaren. (2015) A Comprehensive, Automatically Updated Fungal ITS Sequence Dataset for Reference-Based Chimera Control in Environmental Sequencing Efforts. Microbes and Environment, 30, 145–150.

Oksanen, J., F. Blanchet, G., Friendly, M., Kindt, R., Legendre, P., McGlinn, D., Minchin, P.R., R. B. O’Hara, Simpson, G.L., Solymos, P., M. Stevens, H.H., Szoecs, E., and Wagner, H. (2017). vegan: Community Ecology Package. R package version 2.4-3. https://CRAN.R-project.org/package=vegan.

Öpik, M., Moora, M., Zobel, M., Saks, U., Wheatley, R., Wright, F., Daniell, T., 2008. High diversity of arbuscular mycorrhizal fungi in a boreal herb-rich coniferous forest. New Phytologist, 179, 867–876.

Öpik, M., Vanatoa, a., Vanatoa, E., Moora, M., Davison, J., Kalwij, J.M., Reier, Ü. & Zobel, M. (2010) The online database MaarjAM reveals global and ecosystemic distribution patterns in arbuscular mycorrhizal fungi (Glomeromycota). New Phytologist, 188, 223–241.

Ovaskainen, O., Tikhonoc, G., Norberg, A., Blanchet, F., Duan, L., Dunson, D., Roslin, T. & Abrego, N. (2017) How to make more out of community data? A conceptual framework and its implementation as models and software. Ecology Letters, 20, 1–16.

Paulson, J.N., Stine, O.C., Bravo, H.C. & Pop, M. (2013) Differential abundance analysis for microbial marker-gene surveys. Nature methods, 10, 1200–2.

Peay, K.G. (2014) Back to the future□: natural history and the way forward in modern fungal ecology. Fungal Ecology, 12, 4–9.

Powell, J.R., Parrent, J.L., Hart, M.M., Klironomos, J.N., Rillig, M.C. & Maherali, H. (2009) Phylogenetic trait conservatism and the evolution of functional trade-offs in arbuscular mycorrhizal fungi. Proceedings of the Royal Society B, 4237–4245.

R Core Team (2017). R: A language and environment for statistical computing. R Foundation for Statistical Computing, Vienna, Austria. URL https://www.R-project.org/.

Rao, L.E. & Allen, E.B. (2010) Combined effects of precipitation and nitrogen deposition on native and invasive winter annual production in California deserts. Oecologia, 162, 1035–1046.

Reinhart, K.O. & Callaway, R.M. (2006) Soil biota and invasive plants. New Phytologist, 170, 445–457.

Reinhart, K.O., Tytgat, T., van der Putten, W.H. & Clay, K. (2010) Virulence of soil-borne pathogens and invasion by Prunus serotina. New Phytologist, 186, 484–495.

Richardson, D.M., Allsopp, N., D’Antonio, C.M., Milton, S.J. & Rejmánek, M. (2000) Plant invasions--the role of mutualisms. Biological Reviews, 75, 65–93.

Rognes, T., Flouri, T., Nichols, B., Quince, C. & Mahé, F. (2016) VSEARCH□: a versatile open source tool for metagenomics., 1–22.

Rohland, N. & Reich, D. (2012) Cost-effective, high-throughput DNA sequencing libraries for multiplexed target capture. Genome Research, 22, 939–946.

Rubin, B. E., J. G. Sanders, J. Hampton-Marcell, S. M. Owens, J. A. Gilbert, & C. S. Moreau. 2014. DNA extraction protocols cause differences in 16S rRNA amplicon sequencing efficiency but not in community profile composition or structure. Microbiologyopen 3, 910–921.

Selosse, M.-A., Schneider-Maunoury, L. & Martos, F. (2018) Commentary Time to re-think fungal ecology□? Fungal ecological niches are often prejudged. New Phytologist, 213, 968–972.

Sikes, B.A., Cottenie, K. & Klironomos, J.N. (2009) Plant and fungal identity determines pathogen protection of plant roots by arbuscular mycorrhizas. Journal of Ecology, 97, 1274–1280.

Sikes, B.A., Powell, J.R. & Rillig, M.C. (2010) Deciphering the relative contributions of multiple functions within plant–microbe symbioses. Ecology, 91, 1591–1597.

Taylor, D. L., Walters, W. A., Lennon, N. J., Bochicchio, J., Krohn, A., Caporaso, J. G., & Pennanen, T. (2016) Accurate Estimation of Fungal Diversity and Abundance through Improved Lineage-Specific Primers Optimized for Illumina Amplicon Sequencing. Applied and Environmental Microbiology, 82, 7217–7226.

Treseder, K.A., Allen, E. B., Egerton-Warburton, L. M., Hart, M. M., Klironomos, J. N., Maherali, H. & Tedersoo, L. (2018). Arbuscular mycorrhizal fungi might mediate ecosystem responses to nitrogen deposition: A trait-based predictive framework. Journal of Ecology, 106, 480–489.

Treseder, K.K. & Allen, M.F. (2002) Direct nitrogen and phosphorus limitation of arbuscular mycorrhizal fungi: a model and field test. New Phytologist, 155, 507–515.

Valliere, J.M., Irvine, I.C., Santiago, L. & Allen, E.B. (2017) High N, dry□: Experimental nitrogen deposition exacerbates native shrub loss and nonnative plant invasion during extreme drought. Global Change Biology, 23, 1–13.

Varela-Cervero, S., López-García, Á. & Barea, J.M. (2016a) Spring to autumn changes in the arbuscular mycorrhizal fungal community composition in the different propagule types associated to a Mediterranean shrubland. Plant and Soil, 107–120.

Varela-Cervero, S., López-García, Á., Barea, J.M. & Azcón-Aguilar, C. (2016b) Differences in the composition of arbuscular mycorrhizal fungal communities promoted by different propagule forms from a Mediterranean shrubland. Mycorrhiza, 489–496.

Varela-Cervero, S., Vasar, M., Davison, J., Barea, J.M., Öpik, M. & Azcón-Aguilar, C. (2015) The composition of arbuscular mycorrhizal fungal communities differs among the roots, spores and extraradical mycelia associated with five Mediterranean plant species, Environmental Microbiology, 17, 2882–2895.

Venables, W. N. & Ripley, B. D. (2002) Modern Applied Statistics with S. Fourth Edition. Springer, New York. ISBN 0-387-95457-0/

Vogelsang, K. & Bever, J.D. (2009) Mycorrhizal densities decline in association with nonnative plants and contribute to plant invasion. Ecology, 90, 399–407.

Weber, S.E., Diez, J.M., Andrews, L.V., Goulden, M.L., Aronson, E.L., Allen, M.F., Responses of arbuscular mycorrhizal fungi to multiple coinciding global change drivers. (2018) Manuscript under revision.

Wickham, H. ggplot2: Elegant Graphics for Data Analysis. Springer-Verlag New York, 2009.

Wolfe, B.E. & Klironomos, J.N. (2005) Breaking new ground: Soil communities and exotic plant invasion. Bioscience, 55, 477–487.

